# Phage therapy candidates from Sphae: An automated toolkit for predicting sequencing data

**DOI:** 10.1101/2024.11.18.624194

**Authors:** Bhavya Papudeshi, Michael J. Roach, Vijini Mallawaarachchi, George Bouras, Susanna R. Grigson, Sarah K. Giles, Clarice M. Harker, Abbey L. K. Hutton, Anita Tarasenko, Laura K. Inglis, Alejandro A. Vega, Cole Souza, Lance Boling, Hamza Hajama, Ana Georgina Cobián Güemes, Anca M. Segall, Elizabeth A. Dinsdale, Robert A. Edwards

**Affiliations:** Flinders Accelerator for Microbiome Exploration, College of Science and Engineering, Flinders University, Adelaide, SA, 5042, Australia; Flinders Health and Medical Research Institute, College of Medicine and Public Health, Flinders University, Adelaide, SA, 5042, Australia; Adelaide Medical School, Faculty of Health and Medical Sciences, The University of Adelaide, Adelaide, SA, 5005, Australia; The Department of Surgery - Otolaryngology Head and Neck Surgery, University of Adelaide and the Basil Hetzel Institute for Translational Health Research, Central Adelaide Local Health Network, South Australia, Australia; Department of Biology, San Diego State University, 5500 Campanile Drive, San Diego, CA, 92182, USA; David Geffen School of Medicine, University of California Los Angeles, Los Angeles, CA, USA; Department of Pathology, University of San Diego, 500 Gilman Drive, MC 0612, La Jolla, San Diego, CA, 92093-0612, USA

**Keywords:** phage therapy, antimicrobial resistance, automated pipeline, genome characterization, snakemake

## Abstract

**Motivation:** Phage therapy is a viable alternative for treating bacterial infections amidst the escalating threat of antimicrobial resistance. However, the therapeutic success of phage therapy depends on selecting safe and effective phage candidates. While experimental methods focus on isolating phages and determining their lifecycle and host range, comprehensive genomic screening is critical to identify markers that indicate potential risks, such as toxins, antimicrobial resistance, or temperate lifecycle traits. These analyses are often labor-intensive and time-consuming, limiting the rapid deployment of phage in clinical settings.

**Results:** We developed Sphae, an automated bioinformatics pipeline designed to streamline therapeutic potential of a phage in under ten minutes. Using Snakemake workflow manager, Sphae integrates tools for quality control, assembly, genome assessment, and annotation tailored specifically for phage biology. Sphae automates the detection of key genomic markers, including virulence factors, antimicrobial resistance genes, and lysogeny indicators like integrase, recombinase, and transposase, which could preclude therapeutic use. Benchmarked on 65 phage sequences, 28 phage samples showed therapeutic potential, 8 failed during assembly due to low sequencing depth, 22 samples included prophage or virulent markers, and the remaining 23 samples included multiple phage genomes per sample. This workflow outputs a comprehensive report, enabling rapid assessment of phage safety and suitability for phage therapy under these criteria. Sphae is scalable, portable, facilitating efficient deployment across most high-performance computing (HPC) and cloud platforms, expediting the genomic evaluation process.

**Availability:** Sphae is source code and freely available at https://github.com/linsalrob/sphae, with installation supported on Conda, PyPi, Docker containers.

## Introduction

With the escalating global challenge of antimicrobial resistance comes an increasing demand for alternative treatments against bacterial infections. Bacteriophages, or phages, are viruses that infect bacteria and are ubiquitous in the environment. The use of phages to treat bacterial infections is being explored worldwide as a replacement for antimicrobials. In the US, Australia and parts of Europe, this treatment option is typically administered as a last resort care for severely ill patients under compassionate use [1,2]. For phage therapy to be most effective, thorough safety assessments of the phage isolates must be performed before treatment. This includes experimental testing to confirm that the phage is a pure isolate and can infect the targeted pathogen variant. Additionally, phages are screened to specifically select lytic phages that infect, replicate, and quickly kill the bacterial host over temperate or lysogenic phages that integrate into the host genome during infection and remain stable [3,4]. Temperate phages are not preferred as they can protect the host by improving their fitness and may confer phage resistance through repressor-mediated immunity and/or superinfection exclusion [4,5]. Additionally, phages are screened for large burst sizes and short latent periods to ensure quick and sustained infectivity and high adsorption rates to ensure effectiveness at low concentrations. The presence of these qualities is essential for high virulence to overwhelm the bacteria quickly [6].

Phages and bacteria are locked in an evolutionary arms race where bacterial defense mechanisms like CRISPR-Cas systems co-evolve with phage countermeasures and can propagate throughout bacterial populations [7–9]. Interestingly, it has been shown that the development of phage resistance by the host often coincides with a loss of antibiotic resistance [10], allowing antibiotics to augment phage therapy by eliminating bacteria as they switch from an antibiotic-to a phage-resistant state. This synergy can be enhanced by using phage cocktails consisting of a range of phages with a combined specificity for a broad host range to further reduce the evolution of phage resistance within a bacterial infection. Especially if the cocktail includes phages with distinct mechanisms of host recognition and/or host factors so that resistance to one phage does not confer resistance to all phages [11–13]. Consequently, phage therapy has significant potential to be an effective treatment strategy for combating antibiotic resistance.

Efforts have been renewed to isolate phages for antibiotic-resistant bacterial pathogens in Europe, the US, and Australia. The use of bacteriophages as therapeutic applications is subject to stringent regulatory oversight, particularly concerning toxin production and antimicrobial resistance genes. Ideally, phage isolates are sequenced during screening to predict their genetic potential for safety and efficacy [14–17]. Bioinformatics analysis is now an indispensable component of this approach, ensuring sequencing data is processed efficiently to guide decision-making. For time-sensitive applications, rapid and scalable computational tools are essential, especially for large-scale screening initiatives. However, current analysis workflows can be time-consuming and require manual intervention, limiting their throughput and scalability.

Phage genomes are typically small, with a median size of about 40kb, and can usually be assembled easily into complete genomes. However, the assembly process using default assembly tools obfuscates genome termini signals [15]. The recently published Phables algorithm [18] uses the assembly graph and read coverage to identify and correctly resolve genome termini. Alternatively, the HYbrid and Poly-polish Phage Assembly (HYPPA) method utilizes long-read assemblies in combination with short-read sequencing [19]. Phage genome sequences can also be contaminated with contigs from the bacterial host due to contamination during DNA extraction or due to induction of host prophages, resulting in mixed phage lysates [16]. Tools such as ViralVerify [20] identify and remove putative host contigs [20]. Additionally, phage assemblies may be split over multiple contigs. Therefore, it is important to utilize tools such as CheckV [21] to determine if the assembly represents a single complete phage genome, and in identification of direct terminal repeats. In some cases, even a single phage lysate can yield multiple phage genomes, making such tools indispensable for accurate phage identification [22].

Once assembled, genome annotation tools like Pharokka [23] predict genes and assign biological functions using database searches against genes with known functions. However, assigning biological functions remains challenging, as 65% of viral proteins lack sequence homology to a protein with a known function [24]. Nonetheless, specific genes that serve as markers for temperate lifestyle (such as integrase genes) or confer phage resistance, including a search for toxin, virulence factors, or antimicrobial resistance, are screened for. Such genes are attributed to the risk of horizontal gene transfer (HGT) and propagation of resistance through bacterial populations. These genes are exclusionary criteria for phage therapeutic use, however in cases where lytic phages are unavailable, engineered phages with disabled integrase and repressor functions have been demonstrated as an option [25,26]. Meanwhile, anti-CRIPSR (*Acr*) proteins against their host and depolymerase genes are preferred as they can be advantageous in infection [15]. However, running all these tools sequentially is time-consuming and resource intensive.

Previous studies describe step-by-step tutorials and guidelines for assembling high-quality phage genomes and best practices for predicting and annotating their genes [15,27,28]. We have developed Sphae, a rapid phage characterization workflow designed to streamline the selection of phage therapy candidates. This name is derived from “spae” which means “to foretell” with a modified spelling (s-ph-ae) denoting its specific focus on predicting a phage’s suitability for therapeutic use. This workflow helps quickly select phage therapy candidates based on their genomic potential, which can lead to faster medical interventions and improved patient survival outcomes. We developed this workflow to ensure reproducibility and consistency in the outputs, as using different databases and software versions can influence the results. This workflow is easy to install and run and generates a final summary text file with phage characteristics that anyone can examine to determine the therapeutic potential of a phage.

## Methods

### Workflow input

Sphae requires sequencing reads in fastq format, either paired-end short reads from Illumina or MGI sequencing platforms or unpaired long reads from Oxford Nanopore sequencing platforms. Oxford Nanopore raw sequencing output is in fast5 or pod5 format, which must be basecalled using Guppy (https://community.nanoporetech.com) or Dorado (https://github.com/nanoporetech/dorado) to convert the reads to fastq format before running this workflow.

### Snakemake workflow manager

We utilized the Snakemake workflow manager [29], which facilitates the automated installation of packages and dependencies. We also utilized Snaketool, which provides a user-friendly command line interface for Sphae to make running the pipeline as easy as possible [30] (https://github.com/beardymcjohnface/Snaketool). Workflow managers such as Snakemake provide scalability, reproducibility, reentrancy [31], parallel processing of multiple samples, and integration for running commands and various steps on high-performance computing (HPC) systems and cloud-based environments [30]. Therefore, we employed this template to leverage the capabilities of the Snakemake workflow manager in developing our pipeline for carrying out quality control, genome assembly, and annotation.

### Steps in workflow

1. *Quality control*: Fastp [32] and Filtlong [33] are run to remove low-quality reads and trim adaptor sequences to ensure only high-quality reads are retained for downstream analysis.
2. *Read subsampling*: Rasusa [34] is run to subsample up to 10 million base pairs per sample to keep an ideal 25x to 100x genome coverage for phage assembly [27].
3. *Assembly process*: Paired-end short reads are assembled using MEGAHIT [35], while long-read assemblies are conducted using Flye [36]. Although recent advances in Nanopore sequencing chemistry have reduced the need for long-read polishing [37], medaka [20] is used to correct older, more error-prone reads.
4. *Completeness assessment*: Assembled contigs are classified using:

1. ○ ViralVerify [20] to identify viral, plasmid, or bacteria origin using gene content,
2. ○ CheckV [21] to determine the completeness of the viral contigs by comparing the genomes against a database of viral genomes and identifying the conserved gene markers and regions,
3. ○ and a custom Python script to assess contig connectivity within the assembly graph [18].
4. ○ Overall, only contigs classified as viral by ViralVerify (longer than 1,000 base pairs and having a completeness score of over 70%) are selected for further analysis. In cases of multiple genomes in a sample, each genome is saved as a separate phage genome.
5. *Gene annotation* is performed using Pharokka [23]. Gene prediction is conducted using Phanotate [38] or Pyrodigal [39], followed by functional annotation through comparison with the PHROGs database [40]. In addition, genes are also run against:

1. ○ antimicrobial resistance gene databases: CARD [41],
2. ○ virulence factor database; VFDB [42],
3. ○ CRISPR recognition tool; MinCED [43],
4. ○ Anti CRISPR (Acr) gene detection using AcrDB [44],
5. ○ anti-phage systems using DefenseFinder [45],
6. ○ tRNA genes using tRNAscanSE [46] and tmRNA; ARAGORN [47].
6. Taxonomic assignment is performed within Pharokka,via MASH [48] that compares the genome against the phage INPHARED database [49].
7. *Hypothetical gene analysis*: To address the prevalence of remaining hypothetical genes, Sphae uses:

1. ○ Phold (https://github.com/gbouras13/phold) applies the ProstT5 [50] protein language model to generate a structural representation for each gene. These are compared against a database of predicted phage protein structures using FoldSeek [51] to assign potential functions.
2. ○ The resulting Genbank files are further processed through Phynteny [52], which utilizes a long short-term memory model trained with phage synteny to refine gene function predictions.
8. *Phage therapy suitability*: The annotated genome is systematically analyzed for key markers, including integrase, recombinase, transposase, toxins, antimicrobial resistance, and virulence genes.

### Workflow output

Each workflow step yields a set of files, not all directly pertinent for deciding the therapeutic potential of the phage. Sphae workflow produces a “FINAL” directory containing essential summary files to streamline the output. These files include:

- assembled phage genome (.fasta)
- phage annotations (.gbk)
- genome plot (.png)
- summary table (.tsv): annotations from the three tools, tracking which tool assigned a function to the gene
- summary (.txt): phage characteristics described in Table 1

### Phage sampling and sequencing

*Escherichia coli* strain CoGEN001851 (BEI Resources: Catalog number NR-4359) was received as a glycerol stock from BEI resources. The strain was plated on Brain-Heart Infusion media, supplemented with 1.5% agar (w/v), MgSO_4,_ and MgCl_2_ to a final concentration of 10 mM and 2 mM, respectively. The culture plates were incubated at 37°C for 24 h. The phages were isolated from untreated sewage water (influent) collected from the waste treatment plant in Cardiff, California, as described in [55]. An isolated plaque was selected from each plate and purified further. Phage DNA was then extracted, and *E.coli* phages were sequenced using Oxford Nanopore MinION sequencing according to the manufacturer’s instructions, using Oxford Nanopore Rapid Barcoding Sequencing Kit (SQK-RBK0004) and sequenced on Flowcell R9.4.1 (FLO-MIN106) as described in [55]. The sequencing data were deposited to the Sequence Read Archive (SRA) in Bioproject PRJNA737576. The resulting fast5 reads were run through Guppy v6.0.1 with model dna_r9.4.1_450bps_hac for the Nanopore sequenced isolates. The resulting fastq reads were then run through the Sphae workflow.

### Datasets

The workflow was tested on phages isolated from the above commercially available *E. coli* strains, and with publicly available sequence reads or genomes for *Klebsiella, Salmonella,* and *Achromobacter* phages (Table 2 and S1). Additionally, we included one dataset with five samples that included mixed *Caudovirictes* phages from multiple bacterial hosts to demonstrate the potential of Sphae workflow in assembling and separating each phage (Table 2 and S1). The reads were downloaded from SRA using sra-tools (https://github.com/ncbi/sra-tools) in fastq format as input for Sphae.

### Benchmarking

We benchmarked Sphae’s performance on 5 datasets with 65 samples (Table 2) to compare its functionality and performance. Previous studies have described guidelines [15,27,28] for assembling high-quality phage genomes and annotating their genes; we have used these tutorials as a framework to develop Sphae. All programs and dependency versions used for benchmarking can be found in Table S2. This adaptable workflow is designed with versatility, making it compatible with future updates and new software. As there are no comparable workflows, we assessed the workflow performance using datasets with varying complexities, different numbers of samples, and different sequencing platforms, including samples with single or multiple phages.

Running the workflow in parallel mode processes each phage genome as an individual job, thus speeding the output time. This can be set up on high-performance computing systems (HPC) using a user-provided profile.

### Runtime performance comparison

To evaluate Sphae’s runtime, we measured the wall-clock runtime on a RedHat Linux release 8.10 machine with an AMD EPYC 7551 CPU @ 2.55 GHz. We analyzed sequencing data for a *Klebsiella* phage Amrap using both paired-end and long-read sequencing methods with default settings in Sphae. The analysis was conducted on 6 or 8 threads and 32GB of memory to mimic commonly available consumer hardware. Each paired-end, long-read with polishing, long-read without polishing, and annotate modes were executed 5 times with the same command, and the median wall-clock times with 8 and 16 threads were recorded.

## Results

### Determining complete genomes from assembly

Depending on the complexity and genome coverage of the phage, assembly steps can result in different results (Fig 2). Ideally, the phage genomes are completely assembled into circular or linear genomes (Fig 2A and 2B). In other cases, the direct terminal repeat (DTR) that plays a role in packaging cannot be resolved due to its low complexity during assembly; in this case, the code considers the longer contig as a final genome representation (Fig 2C). Similarly, the DTR regions can cause multiple genomes to be tangled in an assembly graph (Fig 2D). In this case, all the contigs identified as complete phage genomes by CheckV are considered separate phage genomes from a sample. In the final case, the assembly generates fragmented phage genomes; if the contigs are long enough to determine if they are components of a phage genome (Fig 2E), or they may be too fragmented, making it challenging to determine if they are viral (Fig 2F). In both the latter cases, the poor quality of the assembly can lead to poor annotation and, therefore, they are not considered further in the workflow.

### Assembly summary

We assembled 65 samples across the 5 datasets, described in Table 1, using Sphae v1.4.3 with the tools and their version listed in Table S2, which assembled 84 phages. In the summary output (Table S3), we indicate if the assemblies have failed, if the assembly itself has not produced contigs, or if the assembled contigs were fragmented.

In the *E.coli* dataset, some sequences lacked sufficient genome coverage, resulting in unassembled phage genomes (Fig S1). Seven of the 14 samples were assembled, four generated fragmented assemblies, and three failed during assembly (Fig S1). This dataset highlighted how Sphae captures the presence of poorly sequenced samples, suggesting to the user that further sequencing data is required to generate suitable genomes for these phages.

In the case of *Klebsiella* phages, short- and long-read sequences were assembled separately, revealing differences between the two sequencing platforms. Paired-end reads generated complete, circular assemblies with assembly graphs, including one sample featuring one region with multiple contigs tangled together (Fig 2C, Fig S2). Conversely, Nanopore read assemblies resulted in complete, linear phage genomes (Fig 2B, Fig S3), lacking the DTR region (Table S3). With the *Salmonella* and *Achromobacter* phage datasets, complexity arose from samples containing multiple phage genomes. While Sphae was able to assemble phage genomes for each sample (Fig S4, Fig S5), two samples (Se_F6 and Salfasec_13) contained two assembled phage genomes (Fig S4B, Fig S4J), and two samples (Se_F3 and Se_F1) contained three phage genomes (Fig S4C, Fig S4E). This observation aligns with the genome characteristics outlined in the original publication [44], confirming the presence of multiple phages in specific samples. However, three of the 11 samples were potentially contaminated with *E.coli* φX174, likely introduced during the sequencing process. Many Illumina sequences contain φX174 contamination as it is used as a spike-in during 16S rRNA sequencing. Similarly, the *Achromobacter* phage dataset had multiple samples containing two phage genomes per isolate, with 11 out of the 15 phages having either 30Kb, 40Kb, or 70Kb genome lengths. The assembly graph illustrates a structure similar to Fig 2D, with two phages connected by the DTR region (Fig S5).

We further ran Sphae on a dataset comprising five mixed *Caudoviricetes* samples (SRR8788475, SRR8869231, SRR8869234, SRR8869239, and SRR8869241), demonstrating Sphae’s capacity to accurately resolve and separate multiple phages within each sample. For instance, sample SRR8788475 Sphae included four phages, and Sphae assembled all four phages (Fig S6B, Table S3), similarly both phages in SRR8869231were assembled (Fig S6C), three phages from SRR8869239 (Fig S6E) and SRR8869241 (Fig S6F). Interestingly, sample SRR8869234 was listed to include two phages, but Sphae assembled three phages, *Staphylococcus, Klebsiella*, and *Enterobacter* phage. (Fig S6D). Importantly, the resulting assembly graphs across all samples were connected by short sequence fragments (Fig 2D), reflecting the complexity of resolving multiple phages.

### Phage annotation

Phage genes were identified in the 84 assembled phages using PHANOTATE with a default translation table 11. Since phages can potentially use alternative stop codons [57–59], the summary report includes coding density. If low coding density is reported, the assembled phage genomes can be rerun with `sphae annotate`, by changing the config file to utilize Pharokka’s pyrodigal-gv gene prediction [60]. The average coding density for the 84 phages is 95.17%, with a median of 95.20%, confirming the appropriateness of the default translation table.

To enhance the accuracy of gene annotation, Sphae employs an approach that leverages sequence similarity via Pharokka, structural information through Phold, and synteny information through Phynteny. These methods were selected to provide a multi-faceted view of the gene functions and improve annotations. Initially, 8,321 genes were predicted across all 84 phages, with 4,871 genes (58.53%) classified as hypothetical proteins, that includes genes with ambiguous descriptions based solely on sequence similarity searches. However, integrating structural and synteny information, an additional 553 proteins were annotated, highlighting the effectiveness of this combined approach (Fig 3B). Although synteny information did not improve annotation for these datasets, it has been shown to improve annotations in other phages [61]. A summary of the annotated genes based on their PHROG categories showed *Salmonella, Achromobacter,* and mixed phage datasets included genes across all categories, but *E.coli* and *Klebsiella* (8 out of 10) datasets didn’t include genes from transcriptional regulation and integration and excision (Fig 3A).

Figure 3: Functional annotation of phages A) Bubble plot with the proportion of genes annotated to each PHROG functional category across 84 assembled phages. Bubble size indicates the gene proportion per category, and colors differentiate datasets, illustrating the functional diversity. B) Stacked bar plot showing the gene annotation sources: Pharokka (blue), Phold (orange), and remaining hypothetical proteins (green). C) Screening for specific genes that are exclusionary criteria for therapeutic use (integrases, transposase, recombinase, toxin, AMR, and virulence factor genes) and for advantageous genes (anti-CRISPR spacers and defense genes). D) Potential phage therapy candidates were identified based on the absence of exclusionary genes, supporting the selection of safe therapeutic phages.

To identify specific genes of interest for screening these phages for potential therapeutic use, we started with the presence of integrases, which were found in 15 phages from the *Salmonella* dataset, 13 from the *Achromobacter* dataset, and 3 (*Serratia*, *Staphylococcus, Escherichia*) phages from the mixed phage dataset (Figure 3C). The presence of an integrase suggests that these phages are temperate and can persist using the lysogenic cycle. They may protect their host against other phages or express genes altering host functions. Additionally, 10 *Salmonella* phages contained transposase genes, four phages (*Enterobacter,* 2 *Klebsiella, and Staphylococcus*) from the mixed phage dataset contained recombinases, and two (*Klebsiella* and *Escherichia* phage) included two toxin genes. While none of the assembled phage genomes encoded antimicrobial genes, four phages contained virulence factors. Specifically, a phage from the *Salmonella* dataset and three phages (two *Serratia* phages and an *Acinetobacter* phage) from the mixed phage dataset were found to encode immune-modulating virulence genes. While the specific functions of these gene products remain unknown, their presence raises concerns and would disqualify these phages from consideration for therapeutic use. Overall, these 31 phages exhibit markers indicative of a prophage lifestyle or the presence of virulence factors, suggesting they may not be suitable candidates for phage therapy (Figure 4D).

Among the remaining 48 phages, 12 encoded anti-CRISPR proteins: six from *E. coli*, a *Salmonella* phage, and five from *Achromobacter* phages. An *Escherichia* phage from a mixed dataset contained defense genes (Figure 3C). However, 19 of the 48 potential phage therapy candidates came from samples containing multiple phages, necessitating re-isolation to ensure pure cultures. This reduces the viable candidates for phage therapy to 28 phages: 7 against *E. coli*, 19 against *Klebsiella*, 2 against *Achromobacter*, and one against *Pseudomonas* (Figure 3D). No pure candidates were identified from the *Salmonella* dataset.

### Sphae runtime performance

Sphae was executed five times on *Klebsiella* phage Amrap across various sequencing modes, and thread counts to assess differences in median runtime performance. This repetition allowed for robust comparisons, highlighting the variations in efficiency between configurations. Sphae paired-end sequencing mode took a median of 42 minutes on 8 threads but dropped significantly to a median of 9 minutes and 43 seconds on 16 threads. In long-read mode, the workflow was completed in a median of 14 minutes on 8 threads and 7 minutes and 24 seconds on 16 threads. Additionally, when Medaka polishing was omitted during the long-read mode, the median runtime increased to 17 minutes and 9 seconds on 8 threads, but similarly dropped to 8 minutes and 28 seconds on 16 threads. The `sphae annotate` command runs only the annotation steps of the workflow, taking a median of 6 minutes and 13 seconds on 8 threads compared to 6 minutes and 31 seconds on 16 threads (Table S4). Increasing thread count significantly reduces runtime for assembly-related tasks but does not always benefit annotation steps.

## Discussion

Sphae is a reproducible workflow that automates the fundamental bioinformatics steps used in phage therapy to identify candidates for therapeutic use. By integrating 12 bioinformatics tools and nine Python scripts into a unified workflow, Sphae enables seamless execution using a single command. This workflow addresses key challenges in phage therapy by detecting induced prophages, multiple phage species in a sample, direct terminal repeats that could influence horizontal gene transfer. Leveraging Snakemake’s parallelization capabilities, Sphae can process multiple phages simultaneously, often within 10 minutes on 16 threads per phage sample. This makes Sphae a user-friendly solution for clinical applications and allows for rapid detection of phages with therapeutic potential.

We analyzed five datasets including 65 samples, to benchmark Sphae. These datasets included both short-read and long-read sequencing data, assembling 92 phage genomes, of which 28 phages could be used for therapy (Figure 4). We found that phage samples can contain multiple phages, and Sphae reports the characteristics of these phages, making it easier to identify potential candidates for phage therapy that could be further purified if a therapeutic phage candidate is identified. In some instances, contaminants such as *E.coli* φX174 in Illumina sequencing or phage λ in Nanopore sequences were detected as they are used as sequencing controls. In other cases, induced prophages may be present, identifiable by the presence of the same or highly similar sequences across all samples, as demonstrated in the *Achomobacter* dataset in this study. Finally, in cases where the phage fails, Sphae reports at which step the sample failed, if it was during assembly, or if the assembly was fragmented, as demonstrated with the *E.coli* dataset. These findings underscore the importance of thorough characterization and identification of phages for their potential therapeutic use.

### Sphae analysis reveals genomic insights into phage biology

Phage isolation is challenging as a plaque could have multiple phages from the environment, induced prophages, or other contaminants within a single sample. Bacterial isolates frequently contain prophages, and it has been reported that the average prophage density is 2.4 % [62,63]. The prophage excision can contaminate the therapeutic phage lysate, increasing the risk of horizontal gene transfer, including unwanted antimicrobial resistance (AMR) and virulence genes [64,65]. Here, we demonstrate that Sphae effectively captures the prophage contamination cases and informs the user when the sample might require further purification, as shown with the *Achromobacter* dataset, allowing for detecting and excluding phages that could be therapeutically problematic.

Sphae not only assembles and annotates phage genomes from various bacterial hosts but also identifies integrases, transposases, and recombinases - key enzymes involved in the integration and recombination of phage and bacterial DNA [27]. These enzymes are central to horizontal gene transfer (HGT), particularly in facilitating the movement of genes between phages and hosts, which has implications for phage therapy. In the 92 phages analyzed, integrases were detected in 17 phages, transposases in 10 phages, and recombinases in four phages (Fig 3C). While these three genes are associated with temperate lifecycle, recombinases are also part of recombination systems within lytic phages to help with DNA repair and enable the formation of concatemers in genome packaging. Therefore, the presence of recombinase genes is not a clear indication of a temperate lifecycle, further investigation is required.

Another critical aspect of phage biology is phage genome packaging, Phage packaging mechanisms, such as *cos* and *pac* packaging, can influence the likelihood of HGT events [66,67]. For instance, *cos* site phages are less likely to carry out generalized transduction, while *pac* site or headful packaging wherein the bacterial DNA is mistakenly packaged into the phage capsids, facilitating gene transfer between the bacteria [66]. Sphae addresses this by identifying the direct terminal repeats (DTR) in genomes, typically associated with headful packaging, providing insights into the packaging processes. Sphae detected DTRs in 57 of the 92 phages. However, DTRs were detected in 83.82 % Illumina sequenced phage genomes, while none were detected in Nanopore assemblies. *Klebsiella* dataset included 10 phages on both platforms, and DTRs were detected only on Illumina sequences, as noted in the original publication [19]. This discrepancy underscores the importance of sequencing methods and how they influence the detection of this signal. However, current bioinformatic tools cannot easily differentiate between the different packaging mechanisms or detect the correct copy number of repeats, as this influences completeness, which also depends on the type of phage it is [66].

These mechanisms are relevant to determine if AMR genes and virulence factors in the phage can be transferred to the bacterial hosts or introduced into the bacterial population. Sphae, therefore also searches for AMR genes and virulence factors. In the datasets tested, none of the phages encoded AMR, but four genomes included virulence factors. Overall, identification of these genes and reporting them in the summary file is aimed at making the detection of phage therapy candidates effective. As more phages are sequenced, Sphae could serve as a valuable tool not only for identifying therapy candidates but also for advancing studies on phage evolution and host interaction dynamics.

### Sphae follows FAIR principles

This workflow promotes adherence to the Findable, Accessible, Interoperable, Reusable, and Reproducible (FAIR) principles [68]. While developing this workflow, we addressed a number of challenges generally associated with such workflows. This included creating comprehensive documentation with test datasets and structured output, making it easier to navigate and interpret results. While we provide the users with only pertinent outputs in the “RESULTS” directory, the intermediate files are retained so researchers can adapt their approach to resolve assembly complexities.

In instances of assembly failures, Sphae retains intermediate files that outline the steps where the breakdown occurred. For example, poor assemblies resulting from insufficient genome coverage can prompt more sequencing of the sample, if feasible. Additional adjustments such as altering the subsampled reads or switching to alternative assemblers, could also be considered. Alternative assembler options include: SPAdes [69], which handles a full spectrum of k-mers; Canu [70], which utilizes Overlap-Layout-Consensus assemblers; or hybrid assemblies with tools like Unicycler [71] or Plassembler [72], which may be necessary to resolve assembly complexities. Cases of fragmented assemblies connected in an assembly graph can be resolved using Phables [18]. This ensures that even when complete assemblies are not immediately achievable, researchers can refine their approach to resolve assembly complexities, especially in time-sensitive cases.

Sphae workflow also tracks the versions of the software tools used, enhancing reproducibility. We also emphasize the pre-processing steps to ensure standard execution and minimize human error while providing users with readable errors. The challenges and solutions are presented in Table 3 below.

### Sphae is a modular workflow solution

The tools were chosen based on best practices in phage genome characterization [15,27,28]. The focus was on achieving high accuracy and benchmarking for low runtime results. Workflow managers offer the advantage of isolating each software in its environment [29,30]. This means that as the software is improved or new tools are published, they can be quickly added and replace outdated modules. Additionally, more samples can be added to each dataset, and the workflow will run only the new samples, with previously used tool versions if the conda environments were kept. The complete workflow, along with the individual modules, supports reentrancy, allowing steps to be resumed in case they were interrupted.

In Sphae, we have added the option, `sphae run,` to run the entire workflow beginning with sequencing reads to generate final annotations and a summary report. However, the `sphae annotate` module has been included to allow end-users to run only the annotation steps on pre-assembled phage genomes, leveraging Sphae’s approach to improving the number of annotated genes. This module was added for two reasons: first, the assembled genomes can be re-circularized to start from large terminase subunit (*terL*) or other user-selected genes using tools like Dnaapler [73] and visualized with Clinker [74] or pyGenomeViz (https://github.com/moshi4/pyGenomeViz). Second, phages sometimes reassign stop codons by using alternative genetic codes [57,58,75] end-users can change the config file to run pyrodigal-gv [75] for gene prediction in Pharokka instead of the default PHANOTATE [38]. The need for changing tools can be predicted from the coding density reported in the summary.txt file. Phages generally have high coding density to minimize non-coding regions; low-density coding regions suggest that the annotation tools may have incompletely annotated the phage genome [38].

### Future improvements

The ongoing isolation and analysis of phages continue to enhance our grasp of phage biology, evolution and phage-host interactions. Although short-read platforms have traditionally been used for sequencing most phages, there is a growing adoption of long-read sequencing methods such as Oxford Nanopore and PacBio sequencing. An advantage of long-read sequencing is its ability to detect phage DNA modifications, like methylation [76,77], which may play a role in phage resistance and adaptability to microbial communities. While there are over 2,000 phage sequences available in the SRA from Illumina platforms, fewer than 300 phages have been sequenced using long-read technologies such as PacBio and Nanopore platforms (source: https://www.ncbi.nlm.nih.gov/sra). With the increasing availability of long-read sequencing data and the development of automated tools for identifying methylation in phage genomes with minimal manual intervention, we anticipate the integration of this feature into the workflow as a distinct module. Additionally, alternate codon reassignment, recently identified in phage genomes [39,57,58], is now included in Sphae offering users insights into unique coding adaptions, and insights into coding adaptations relevant to host specificity. Tools like Prfect that predict programmed ribosomal frameshifts producing longer proteins [78], also present an exciting future integration. These enhancements will enable end-users to explore these specialized genome feature, as our understanding of phage biology and evolution improves. The tools and modules within Sphae will be regularly updated to accommodate these advancements to include useful summary reports, ensuring users can easily access and interpret the latest advancements in a user-friendly manner.

### Conclusions

Sphae is a bioinformatics workflow designed to quickly and comprehensively characterize phage isolates and identify phage therapy candidates, addressing the urgent need for effective alternatives to combat antimicrobial resistance. By seamlessly integrating high-quality genomic data and automated analysis, Sphae not only enhances our understanding of phage biology and evolution but also empowers researchers to make informed decisions in the fight against resistant bacterial pathogens.

### Key points

- Sphae optimizes the bioinformatics workflow for phage genome characterization, allowing for the evaluation of potential therapeutic candidates in under ten minutes.
- Sphae is capable of handling both short and long-read sequencing data and generates a comprehensive summary report detailing the presence of key marker genes, including virulence factors, antimicrobial resistance genes, and lysogeny functions (such as integrases, transposases, and recombinases) to assess the suitability of phages for therapy.
- Sphae is designed for scalability and portability, enabling easy deployment on high-performance computing (HPC) and cloud platforms while adhering to FAIR principles.
- A modular architecture allows for the seamless integration of new tools and adaptations

Biographical Note: Sphae is a bioinformatics workflow designed for phage genome characterization, providing a summary report to quickly identify potential phage therapy candidates in clinical settings for time-sensitive cases.

## Funding

This work was supported by an award from NIH NIDDK RC2DK116713 and the Australian Research Council DP220102915.

## Acknowledgments

This research/project was undertaken with the assistance of resources and services from Flinders University using the DeepThought cluster [62], Australian Nectar Research Data Commons (ARDC) Nectar Infrastructure, the Pawsey Supercomputing Research Centre, and the National Computational Infrastructure (NCI), which is supported by the Australian Government.

## Author contributions

B.P. developed the Sphae tool and wrote the manuscript. M.J.R., V.M., G.B., S.R., and L.I. assisted in testing the workflow and made significant contributions to the workflow development. S.K.G., C.H., A.L.K.H., A.T., A.A.V., C.S., L.B, H.H., A.G.C.G., and A.B., were responsible for collecting, isolating, and culturing the phages used to develop and validate this workflow. A.M.S., E.A.D., and R.A.E. conceived the project, provided editorial feedback, and contributed valuable input on key steps to be included in the workflow.

## Completing interests

There are no competing interests

## Data and code availability

The data generated <u>in</u> this article are available in Sequence Read Archive, Bioproject ID: PRJNA737576, other publicly available data was downloaded from SRA, from Bioprojects PRJNA914245, PRJNA914245, PRJEB33638, and PRJNA222858. The Sphae workflow, along with documentation, is available on GitHub at https://github.com/linsalrob/sphae.

## Supplementary Files

Fig S1: (A) Sequencing depth evaluation of E. coli datasets. Samples with high sequencing depth (E.coli_17, E.coli_27, E.coli_29, E.coli_31, E.coli_32, E.coli_34, E.coli_36, and E.coli_37) successfully assembled into complete phage genomes. In contrast, samples with low sequencing depth (E.coli_26, E.coli_28, E.coli_33, E.coli_33_1, E.coli_35, and E.coli_39) produced either no contigs or fragmented contigs during assembly. (B-L) Bandage plots of 10 E. coli phages, showing assembly results for (B) E.coli_17, (C) E.coli_27, (D) E.coli_28, (E) E.coli_29, (F) E.coli_31, (G) E.coli_32, (H) E.coli_33 (fragmented), (I) E.coli_34, (J) E.coli_35 (fragmented), (K) E.coli_36, (L) E.coli_37 (fragmented). Three samples, E.coli_33_1, E.coli_39, and E.coli_26, failed to assemble.

Fig S2: (A) Sequencing depth evaluation of Klebsiella short-read datasets. (B-L) Bandage plot of the 10 phages; each included only one phage per sample, B) Kleb-SR_Whistle, C) Kleb-SR_Amrap, D) Kleb-SR_Emom, E) Kleb-SR_Saitama, F) Kleb-SR_Tokugawa, G) Kleb-SR_Cornelius, H) Kleb-SR_Speegle, I) Kleb-SR_Mera, J) Kleb-SR_Toyotomi, K) Kleb-SR_Oda. The width of the lines in the bandage plots are random and do not reflect genome lengths.

Fig S3: (A) Sequencing depth evaluation of Klebsiella long-read datasets. (B-L) Bandage plot of the 10 phages; each included only one phage per sample, B) Kleb-SR_Whistle, C) Kleb-SR_Amrap, D) Kleb-SR_Emom, E) Kleb-SR_Saitama, F) Kleb-SR_Tokugawa, G) Kleb-SR_Cornelius, H) Kleb-SR_Speegle, I) Kleb-SR_Mera, J) Kleb-SR_Toyotomi, K) Kleb-SR_Oda.

Fig S4: (A) Sequencing depth evaluation of Salmonella short-read datasets. (B-L) Bandage plot of the 11 Salmonella phages with most samples including a single phage, except two samples, (B) SAL_Se_F6 (two phages), (C) SAL_Se_F3 (three phage), (D) SAL_Se_F2, (E) SAL_Se_F1 (three phages), (F) SAL_Se_ML1, (G) SAL_Se_EM4, (H) SAL_Se_EM3, (I) SAL_Se_EM2, (J) SAL_Salfasec_13 (two phages), (K) SAL_Se_EM1, (L) SAL_Se_AO1. The width of the lines in the bandage plots are random and do not reflect genome lengths.

Fig S5. (A) Sequencing depth evaluation of 15 Achromobacter short-read datasets. (B-M) Bandage plots of 12 of the 15 assembled Achromobacter phages: (B) Achrom_Axy06 (one phage), (C) Achrom_Axy09 (two phages), (D) Achrom_Axy24 (two phages), (E) Achrom_Axy23 (two phages), (F) Achrom_Axy10 (two phages), (G) Achrom_Axy12 (one phage), (H) Achrom_Axy13 (two phages), (I) Achrom_Axy21 (two phages), (J) Achrom_Axy16 (one phage), (K) Achrom_Axy19 (two phages), (L) Achrom_Axy18 (two phages), and (M) Achrom_Axy22 (two phages). Three samples are not shown, as their bandage plots were too large for display. Line widths in the bandage plots are arbitrarily scaled and do not represent actual genome lengths.

Fig S6: (A) Sequencing depth evaluation of the five mixed dataset phages. (B-F) Bandage plots, (B) SRR8788475 includes four phages, (C) SRR8869231 includes two, (D) SRR8869234 includes three phages, (E) SRR8869239 includes three phages, (F) SRR8869241 includes three phages.

Table S1: Summary of 65 phage samples from five datasets used to benchmark Sphae performance in this study

Table S2: Software programs and versions utilized in Sphae v1.4.3 for benchmarking and analysis

Table S3: Assembly and annotation results for the 65 phage genomes analyzed in this study Table S4: Runtime Benchmarking of Sphae on *Klebsiella* phage Amrap

---

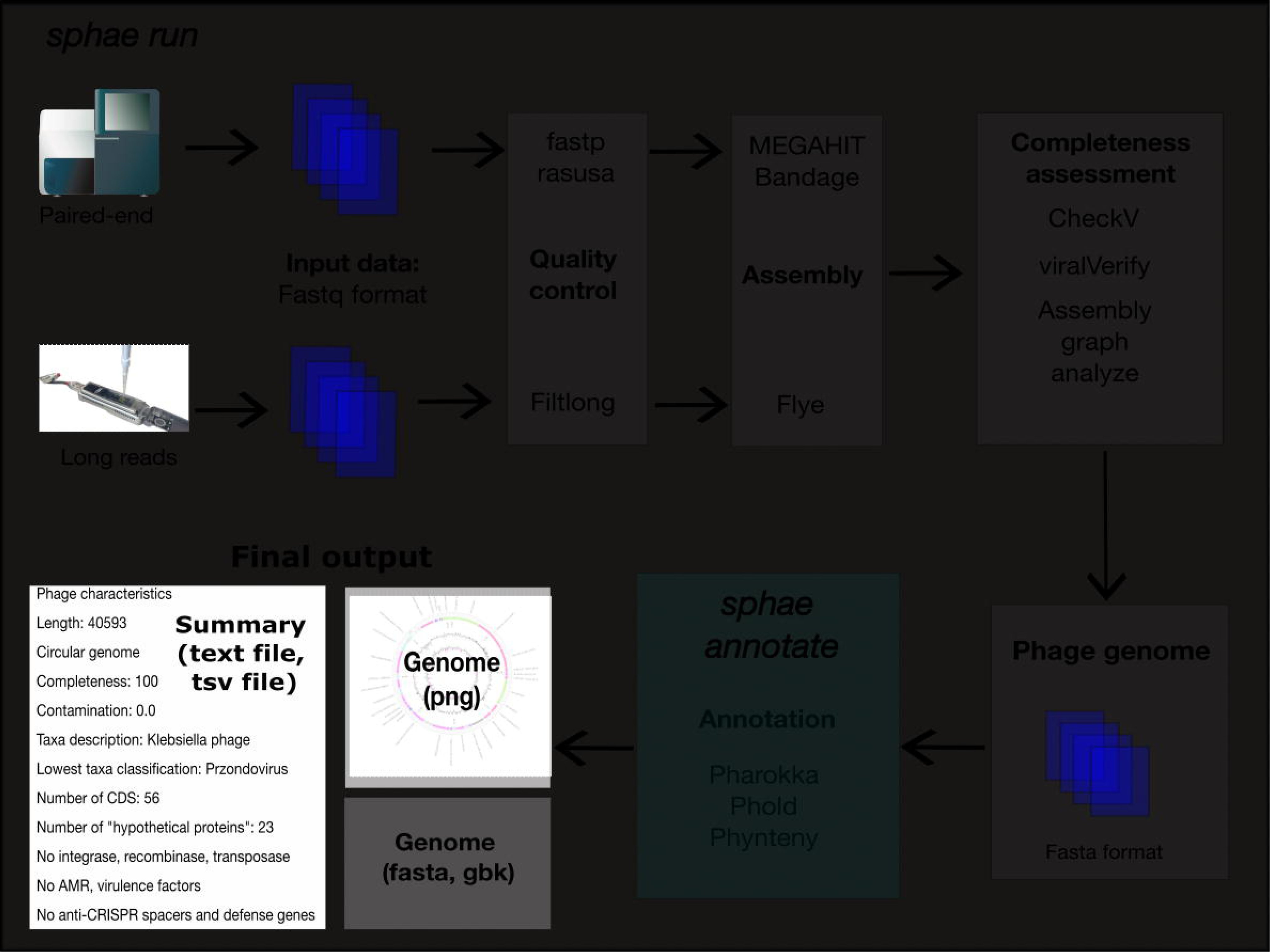

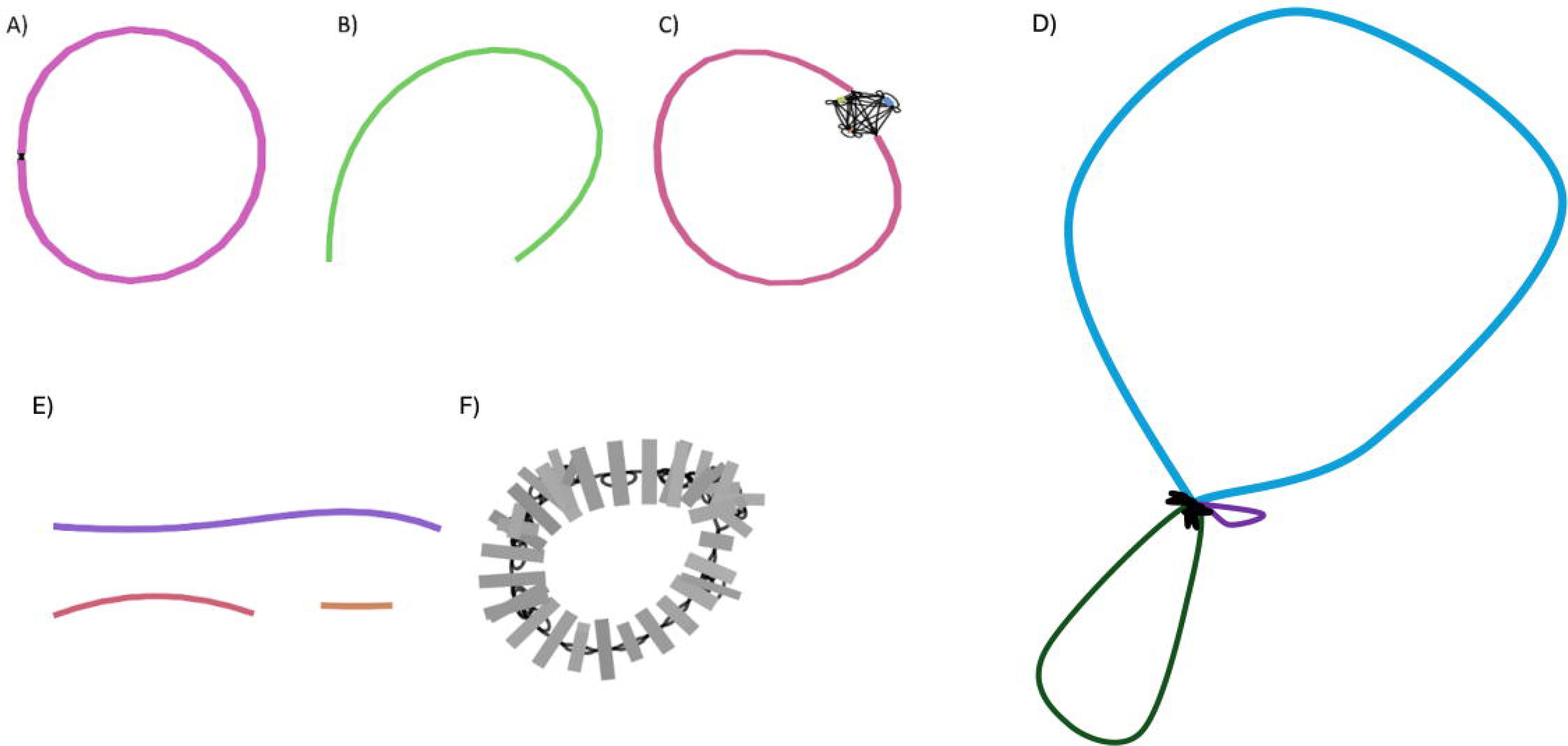

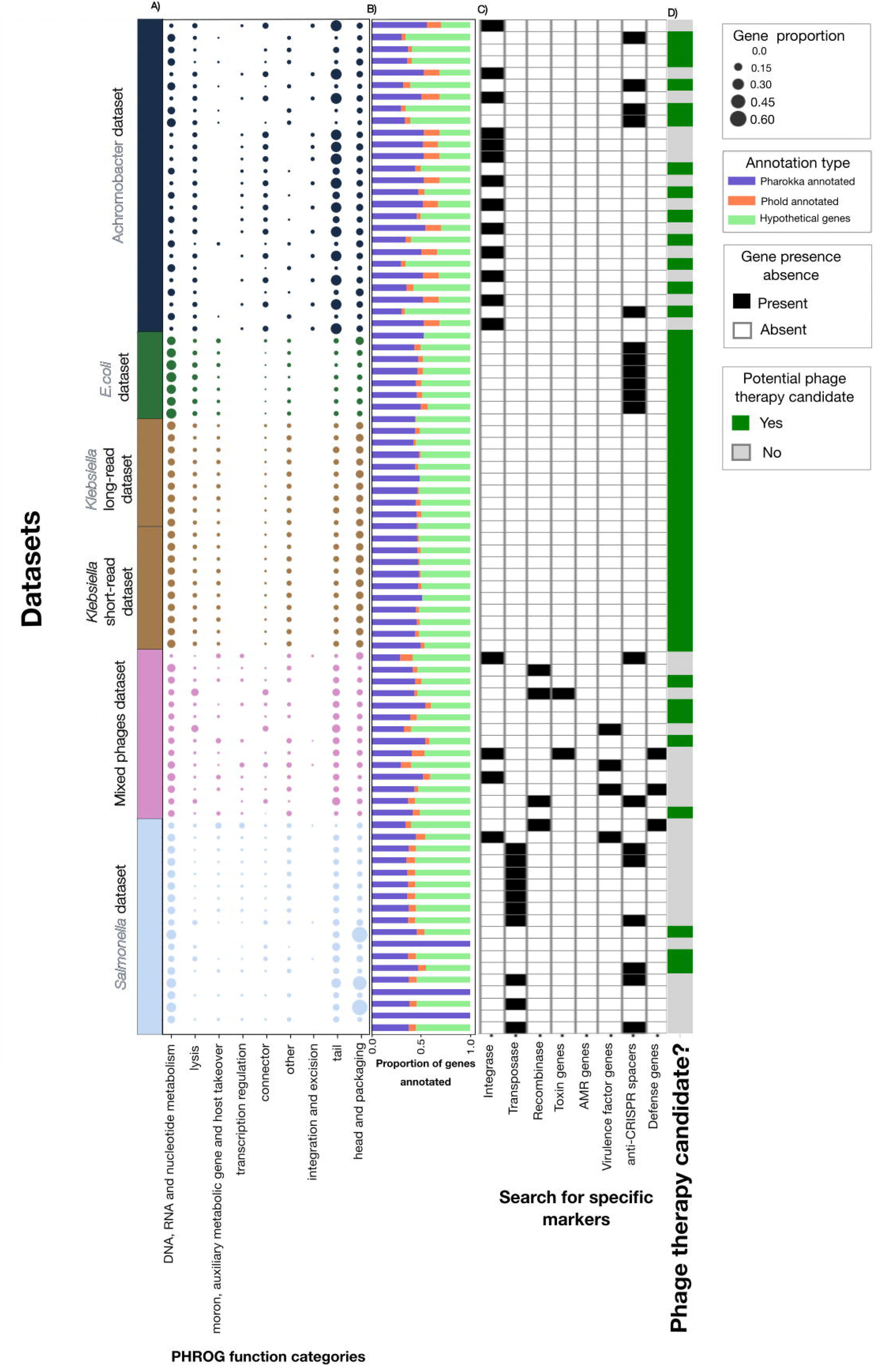

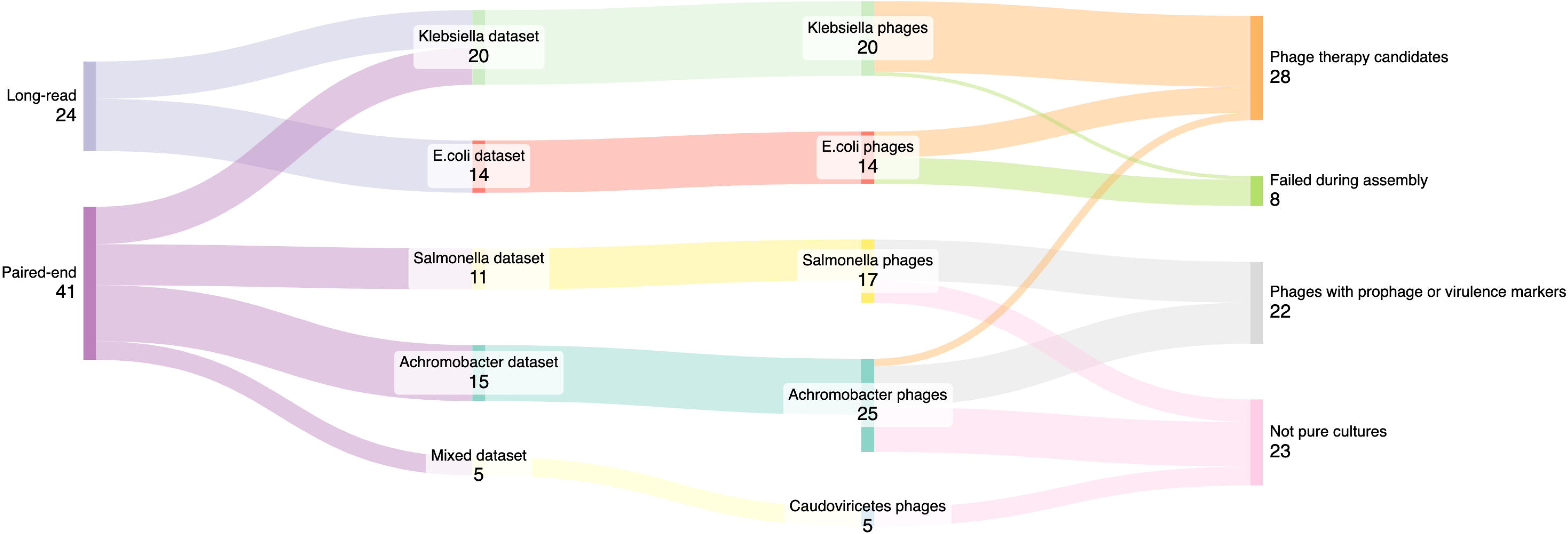

| Challenges | Solution |
| --- | --- |
| Variability in tools and programming languages | Snakemake workflow manager allows the integration of tools written in multiple languages. |
| Lack of version, parameters documentation, and installation of multiple programs | Snakemake allows logging each step, keeping track of the tool version and the command run with the default parameters listed.<br>Each software is automatically downloaded to its separate conda environment with dependencies or via a pre-built Docker/Singularity container. |
| Portability of the workflow | The workflow is available through conda, pip, pre-built containers, and source installation in GitHub or via a pre-built Docker/Singularity container. |
| Hardware and software dependencies | The workflow's configuration file includes resource information that the user can update for the system on which the workflow is running. In addition, a pipeline can be set to talk to job schedulers on high-performance computing (HPC) systems. |
| Error handling | Provide detailed logs with information identifying the step at which the error occurred for each rule and an overall snakemake .log file. |
| Addition of new tools | New tools can be quickly added as a new rule to the workflow. This critical feature allows new and improved tools to be integrated as they are developed. |

| Study | Number of phage samples | Sequencing platform | Bacterial host | Bioproject | Reference |
| --- | --- | --- | --- | --- | --- |
| *E.coli* phages | 14 | MinION | *E.coli* strain CoGEN001851(B<br>EI Resources:<br>Catalog number,<br>NR-4359) | PRJNA737<br>576 | This study |
| *Klebsiella* phages | 20 | MinION<br>Illumina<br>NextSeq | *Klebsiella michiganensis*,<br>*Klebsiella oxytoca*,<br>*Klebsiella quasipneumoniae*,<br>*Klebsiella variicola* | PRJNA914<br>245 | [19] |
| *Salmonella* phages | 11 | Illumina<br>MiSeq | *Salmonella enterica* subsp.<br>*enterica* serovar<br>*Typhimurium*<br>(ATCC 14028S) | PRJNA914<br>245 | [22] |
| *Achromobacter* phages | 15 | Illumina<br>MiSeq | *Achromobacter xylosoxidans*<br>strain 19-32 | PRJEB336<br>38 | [56] |
| Mixed<br>*Caudovirictes* phages | 5 | Illumina<br>MiSeq |  | PRJNA222<br>858 | NA |

| Phage characteristics | Value | Explained |
| --- | --- | --- |
| Sample name | Bc01 | sample name |
| Total length of reads after QC and subsampling | 5,363,156 bp | total length of reads used for assembly to help calculate genome coverage |
| Length | 100743 | length of the phage genome assembled |
| Circular | False | Was the genome assembled to be circular, according to the information provided in the assembly graph? For more information, you can visualize the file ending in .gfa with Bandage [53] |
| Graph connections | 0 | If the assembly generated fragmented contigs due to low coverage, but the graph shows potential connections, offering clues for identifying terminal repeats and low complexity regions. For more information, you can visualize the file ending in .gfa with Bandage [53]. |
| Direct Terminal Repeat (DTR) Found |  | is DTR detected by CheckV [21] in the phage contig |
| Completeness | 100.0 | phage completeness score from CheckV |
| Contamination | 0.0 | contamination score from CheckV |
| Taxon description | *Kehishuvirus* sp. tikkala | assigned taxon name from Pharokka [23] output, comparing the phage genome against the INPHARED database [49] using Mash [48] |
| Taxa result: matching hashes | 972/1000 | How close the phage isolated is to the assigned taxon? The results are from the Pharokka Mash sketch against the INPHARED database |
| Lowest taxon classification | *Kehishuvirus* | the lowest taxon rank assigned |
| Isolation host of the described taxa | *Bacteroides cellulosilyticus* | Bacterial host of the assigned taxa from the INPHARED database |
| Number of CDS | 154 | number of genes identified in the genome from Pharokka result |
| Total number of CDS annotated as 'hypothetical protein' | 91 | counting only the genes annotated as hypothetical, haven't been assigned a biological function or have ambiguous descriptions in Phynteny [52] output |
| GC content (proportion) | 0.35 | GC content from Pharokka result |
| Percent coding density | 91.3 | Phages generally have high coding capacity, so if the density is low, it could be a warning that the gene calling did not work well for this phage |
| Prophage or temperate lifestyle markers | no Integrases,<br>no recombinases,<br>no transposases | These genes indicate the phage can have a temperate lifestyle, which would most likely exclude the phage from use in therapy. The results are from Pharokka, Phold, and Phynteny searches. |
| Toxin genes | no toxins found | Search for genes with the word "toxins" in the gene description from the final Phynteny output. |
| Virulence genes | no antimicrobial resistance (AMR) genes found,<br>no virulence factors found | Search against the CARD [54] and VFDB [42] databases using Pharokka and Phold results. |
| Defense genes | No anti-CRISPR or spacers found<br>No defense genes found | Pharokka and Phold search the genes against ACR [44] and DefenseFinder [45] databases. |

## Notes

### Competing Interest Statement

The authors have declared no competing interest.

