## Supplementary Figures for "Phage therapy candidates from Sphae: An automated toolkit for predicting sequencing data"

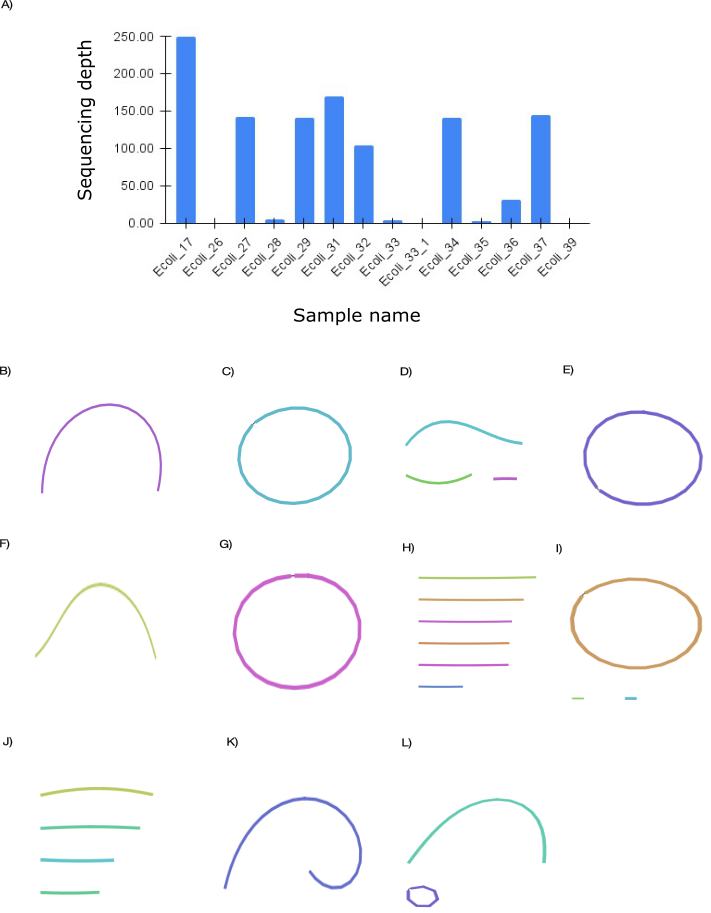

#### Fig S1: (A) Sequencing depth evaluation of *E. coli* datasets. Samples with high sequencing depth (E.coli_17, E.coli_27, E.coli_29, E.coli_31, E.coli_32, E.coli_34, E.coli_36, and E.coli_37) successfully assembled into complete phage genomes. In contrast, samples with low sequencing depth (E.coli_26, E.coli_28, E.coli_33, E.coli_33_1, E.coli_35, and E.coli_39) produced either no contigs or fragmented contigs during assembly. (B-L) Bandage plots of 10 *E. coli* phages, showing assembly results for (B) E.coli_17, (C) E.coli_27, (D) E.coli_28, (E) E.coli_29, (F) E.coli_31, (G) E.coli_32, (H) E.coli_33 (fragmented), (I) E.coli_34, (J) E.coli_35 (fragmented), (K) E.coli_36, (L) E.coli_37 (fragmented). Three samples, E.coli_33_1, E.coli_39, and E.coli_26, failed to assemble.

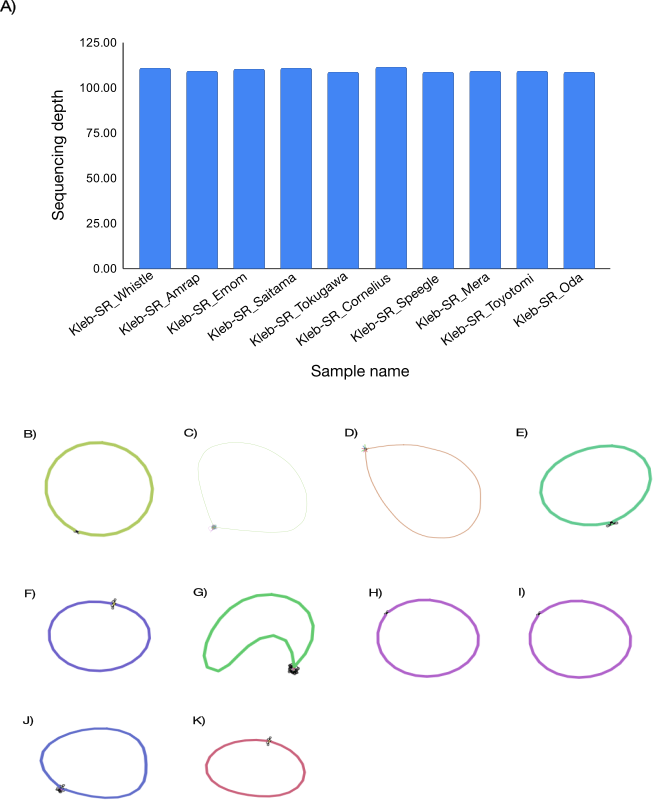

#### Fig S2: (A) Sequencing depth evaluation of *Klebsiella* short-read datasets. (B-L) Bandage plot of the 10 phages; each included only one phage per sample, B) Kleb-SR_Whistle, C) Kleb-SR_Amrap, D) Kleb-SR_Emom, E) Kleb-SR_Saitama, F) Kleb-SR_Tokugawa, G) Kleb-SR_Cornelius, H) Kleb-SR_Speegle, I) Kleb-SR_Mera, J) Kleb-SR_Toyotomi, K) Kleb-SR_Oda. The width of the lines in the bandage plots are random and do not reflect genome lengths.

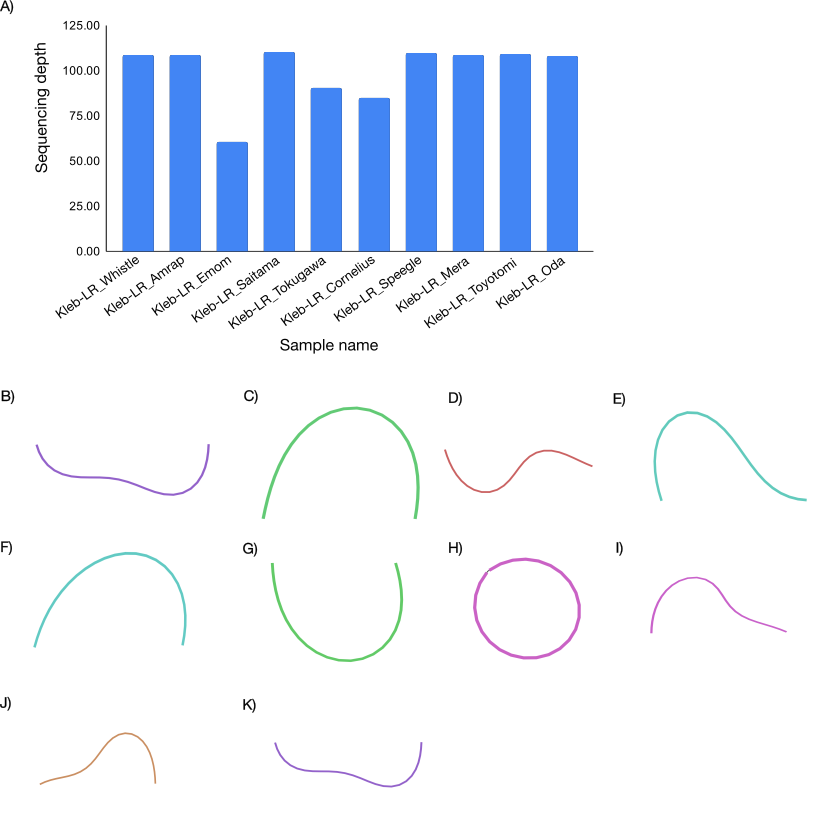

#### Fig S3: (A) Sequencing depth evaluation of *Klebsiella* long-read datasets. (B-L) Bandage plot of the 10 phages; each included only one phage per sample, B) Kleb-SR_Whistle, C) Kleb-SR_Amrap, D) Kleb-SR_Emom, E) Kleb-SR_Saitama, F) Kleb-SR_Tokugawa, G) Kleb-SR_Cornelius, H) Kleb-SR_Speegle, I) Kleb-SR_Mera, J) Kleb-SR_Toyotomi, K) Kleb-SR_Oda.

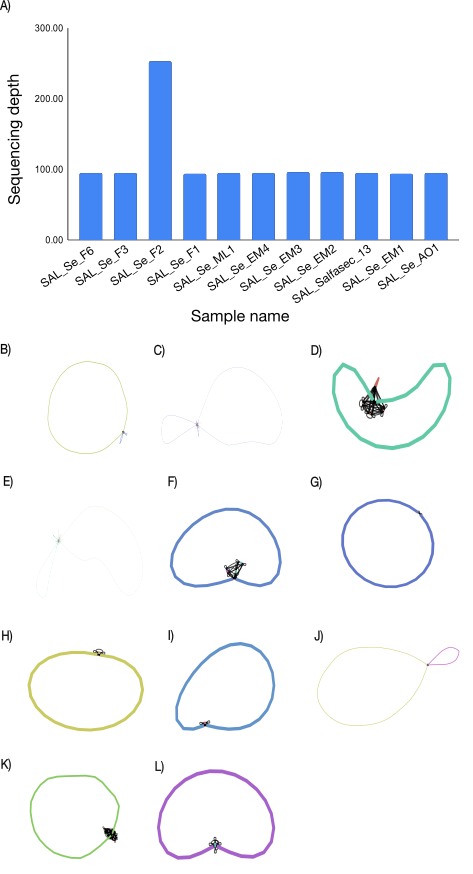

#### Fig S4: (A) Sequencing depth evaluation of *Salmonella* short-read datasets. (B-L) Bandage plot of the 11 Salmonella phages with most samples including a single phage, except two samples, (B) SAL_Se_F6 (two phages), (C) SAL_Se_F3 (three phage), (D) SAL_Se_F2, (E) SAL_Se_F1 (three phages), (F) SAL_Se_ML1, (G) SAL_Se_EM4, (H) SAL_Se_EM3, (I) SAL_Se_EM2, (J) SAL_Salfasec_13 (two phages), (K) SAL_Se_EM1, (L) SAL_Se_AO1. The width of the lines in the bandage plots are random and do not reflect genome lengths.

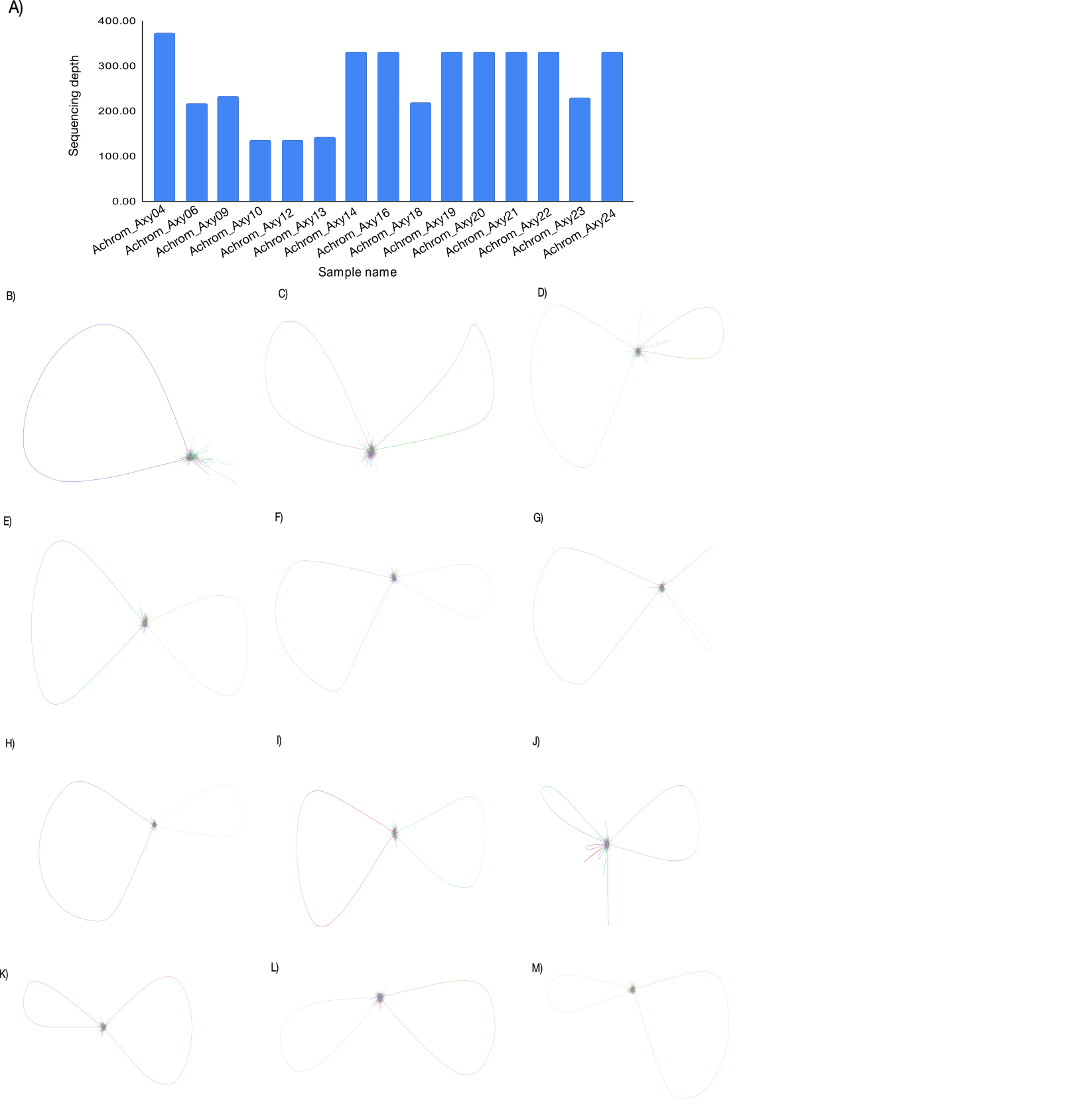

#### Fig S5. (A) Sequencing depth evaluation of 15 Achromobacter short-read datasets. (B-M) Bandage plots of 12 of the 15 assembled Achromobacter phages: (B) Achrom_Axy06 (one phage), (C) Achrom_Axy09 (two phages), (D) Achrom_Axy24 (two phages), (E) Achrom_Axy23 (two phages), (F) Achrom_Axy10 (two phages), (G) Achrom_Axy12 (one phage), (H) Achrom_Axy13 (two phages), (I) Achrom_Axy21 (two phages), (J) Achrom_Axy16 (one phage), (K) Achrom_Axy19 (two phages), (L) Achrom_Axy18 (two phages), and (M) Achrom_Axy22 (two phages). Three samples are not shown, as their bandage plots were too large for display. Line widths in the bandage plots are arbitrarily scaled and do not represent actual genome lengths.

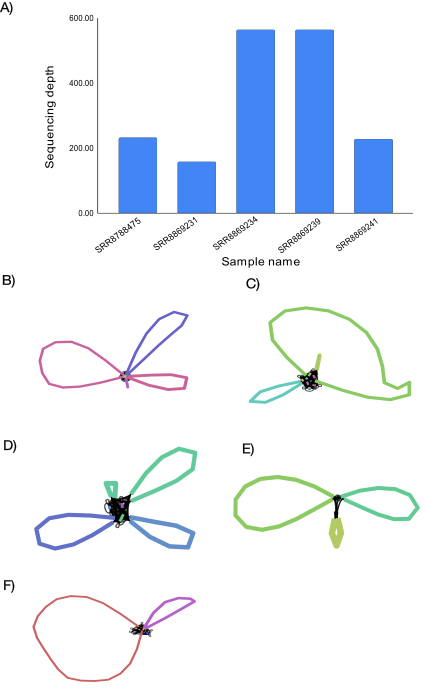

#### Fig S6: (A) Sequencing depth evaluation of the five mixed dataset phages. (B-F) Bandage plots, (B) SRR8788475 includes four phages, (C) SRR8869231 includes two, (D) SRR8869234 includes three phages, (E) SRR8869239 includes three phages, (F) SRR8869241 includes three phages.
